# Re-introduction of dengue virus serotype 2 in the state of Rio de Janeiro after almost a decade of epidemiological silence

**DOI:** 10.1101/661868

**Authors:** Torres María Celeste, Fernanda de Bruycker Nogueira, Fernandes Carlos Augusto, Louzada Silva Meira Guilherme, Ferreira de Aguiar Shirlei, Chieppe Alexandre Otavio, Bispo de Filippis Ana María

**Affiliations:** Laboratório de Flavivírus, Instituto Oswaldo Cruz, Fiocruz, Rio de Janeiro, Brazil; Laboratório Central Noel Nutels – LACEN, Rio de Janeiro, Brazil; Secretaria Estadual de Saúde, Superintendência de Vigilância Sanitária do Estado do Rio de Janeiro, Brazil

## Abstract

The Asian/American genotype of dengue virus serotype 2 (DENV-2) has been introduced in Brazil through the state of Rio de Janeiro around 1990, and since then it has been spreading and evolving, leading to several waves of dengue epidemics throughout the country that cause a major public health problem. Of particular interest has been the epidemic of 2008, whose highest impact was evidenced in the state of Rio de Janeiro, with a higher number of severe cases and mortality rate, compared to previous outbreaks. Interestingly, no circulation of DENV-2 was witnessed in this region during the preceding 9-year period. By early 2010, phylogenetic analysis of the 2008 epidemic strain revealed that the outbreak was caused by a new viral lineage of the Asian/American genotype, which was pointed as responsible for the outbreak severity as well. The same scenario is repeating in 2019 in this state; however, only a few cases have been detected yet. To provide information that helps to the understanding of DENV-2 dynamics in the state of Rio de Janeiro, and thereafter contribute to public health control and prevention actions, we employed phylogenetic studies combined with temporal and dynamics geographical features to determine the origin of the current viral strain. To this effect, we analyzed a region of 1626 nucleotides entailing the Envelope/NS1 viral genes. Our study reveals that the current strain belongs to the same lineage that caused the 2008 outbreak, however, it is phylogenetically distant from any Brazilian strain identified so far. Indeed, it seemed to be originated in Puerto Rico around 2002 and has been introduced into the state in late 2018. Taking into account that no DENV-2 case was reported over the last decade in the state (representing a whole susceptible children generation), and the fact that a new viral strain may be causing current dengue infections, these results will be influential in strengthening dengue surveillance and disease control, mitigating the potential epidemiological consequences of virus spread.

**Author Summary:** By the time dengue virus serotype 2 (DENV-2) was introduced into Brazil through the state of Rio de Janeiro in 1990, the first dengue hemorrhagic cases started to evidence as well. Years of seasonal outbreaks were followed by almost ten years oy epidemiological silence in the state. However, in 2007 this serotype was re-introduced into the state causing one of the worst dengue epidemics ever described in the country. The same viral genotype was involved, however, a different viral lineage was detected and pointed as responsible for the outbreak severity. This same scenario could repeat nowadays in the state of Rio de Janeiro. Since new DENV-2 cases are being detected in this region, we analyzed the identity and origin of the viral strain obtained from two infected patients. Phylogeny combined with temporal and geographical analyses of viral sequences demonstrated that the strain causing 2019’s dengue cases belonged to the same lineage as the one causing the outbreak in 2008, but to a different subgroup, and might have originated in Puerto Rico and entered the state in recent times. These results may represent a crucial starting point for strengthening Brazilian surveillance systems and disease control, helping to reduce the impact of a potential epidemic of great magnitude.

## Introduction

Arboviral infections have been re-emerging in Brazil over the last decades, threatening the country and causing a constant burden for public health [1, 2]. Human travelers and extensive migrations have facilitated the spread of arbovirus potentially threatening for human health through the globe [3 4]. As a tourist magnet, the state of Rio de Janeiro, in southeast Brazil, has been facing large arbovirus epidemics year by year [5].

Dengue virus (DENV) is an arbovirus that belongs to the *Flaviviridae* family, *Flavivirus* genus, and is transmitted to humans by mosquitoes from genera *Aedes*. Four genetically related but antigenically distinct DENV serotypes have been described: DENV-1, DENV-2, DENV-3, and DENV-4. Besides Brazil, DENV is widely distributed across tropical and subtropical areas of the world [6].

In 1986 the serotype DENV-1 was identified in the state of Rio de Janeiro, and since then, dengue fever became a public health burden for Brazil [7]. By early 1990, the Asian/American (As/Am) genotype of serotype DENV-2 was isolated from patients from the metropolitan region of the state [8]. From then on, its spreading gave rise to the epidemic of the years 1990-1991, leading to the advent of the first severe forms of the disease [9]. These two serotypes were also responsible for the epidemic waves of 1995-1996 and 1998, from which the virus spread rapidly to other districts [10]. Already in 2000, the serotype DENV-3 was firstly isolated from a patient in the state of Rio de Janeiro as well. The introduction of this new serotype was associated with an increase in the number of cases during the summer of 2001-2002, causing the largest epidemic so far in the state [11]. This serotype circulated predominantly in the state until 2007, when serotype DENV-2 reemerged, causing a massive and severe epidemic during 2008 that affected the entire country. The state of Rio de Janeiro, however, was the most affected, with more than 235,000 reported cases, over 13,000 hospital admissions and 263 deaths [12], which represented increased disease severity and a higher rate of mortality, compared to previous outbreaks [13]. This epidemic caused staggering human and economic costs, with more than US$1 billion being spent at the national level on dengue prevention and control [14]. Drumond et al. demonstrated that the virus causing the 2008 epidemic belonged indeed, to the same genotype that was introduced around 1990, but to a different lineage [15]. In fact, this new lineage presented genetic differences potentially associated with a more pathogenic clinical outcome of the disease [16]. Henceforth, in 2011, the serotype DENV-4 was identified in the municipality of Niterói, state of Rio de Janeiro, and spread the following year to other districts in the state, accounting for most cases of dengue in 2012 and 2013 [17–19]. The circulation of this serotype decreased after that, giving way again to serotype DENV-1 [20]. During the following years, these two serotypes kept on circulating in the state, however, since late 2014, different arboviruses such as chikungunya virus (CHIKV), Zika virus (ZIKV), and yellow fever virus (YFV), joined alternating their circulation [19]. DENV-2 cases remained undetectable in the state since 2010, until 2017, when one new case was reported in the capital of the state [21]. Consequently, the statal health department claimed to intensify surveillance, even though DENV detection during 2017 was low, and the focus was mostly on surveillance of yellow fever cases and epizootics, and immunization actions as well [22]. However, it was only in 2018, when the Information System for Notifiable Diseases (SINAN) and the Laboratory Sample Management System (GAL) registered three new different DENV-2 cases in the state of Rio de Janeiro [23].

After 2009 however, DENV-2 maintained a basal and low circulation at a national level, always below serotypes DENV-1 and DENV-4. Less than 4% of the notified dengue cases belonged to serotype DENV-2 mainly in the north and northeast regions of the country [18, 19].This scenario abided until 2016, but in 2017, even being the year with the lowest number of registered dengue cases nationally, the central-west region of the country started to show a growing predominance of DENV-2 [24]. During 2018, it kept on spreading at a low rate, obtaining on average, similar figures as the ones from 2017 [25]. However, throughout this current year, DENV-2 cases reached notification rates 282% superior as the preceding year, and its circulation has been already confirmed also in the southeast region of the country, including the state of Rio de Janeiro [26].

Considering that after almost a decade of epidemiological silence of DENV-2 in the state, a whole new generation is naïve to this serotype infection and concerned about a potential new outbreak of great magnitude in the state, we sought to briefly analyze the virus phylogeny and its origin, to contribute to public health policies by providing additional information that may help to strengthen monitoring, control and surveillance measures of this viral strain spreading.

## Materials and methods

### Ethical statement

The samples analyzed in this study belong to the collection from the Laboratory of Flavivirus, IOC/FIOCRUZ, Rio de Janeiro, Brazil, and were obtained through the passive surveillance system from an ongoing Project approved by resolution number CAAE 90249218.6.1001.5248 from the Oswaldo Cruz Foundation Ethical Committee in Research (CEP 2.998.362), Ministry of Health-Brazil. Samples were chosen anonymously, based on the laboratorial results.

### Study samples

Sera samples were obtained from ten patients with suspected dengue fever from the districts of Vassouras, Volta Redonda, Nova Iguaçu e Mangaratiba (west of the state of Rio de Janeiro), between January and March of the current year.

They were sent to the Central Laboratory of Rio de Janeiro LACEN/RJ for diagnostic confirmation through RT-PCR under Lanciotti’s protocol [27]. DENV-2 strain 40247 (Produced by Bio-Manguinhos/FIOCRUZ, RJ) [28] was employed as a positive control, while a normal human serum sample collected from a healthy donor was used as a negative one.

### DENV-2 genetic sequencing

Viral strain sequencing was performed by the Regional Reference Flavivirus Laboratory at Instituto Oswaldo Cruz/FIOCRUZ in Rio de Janeiro. Briefly, viral RNA was extracted from 140 ul of serum samples using the QIAamp Viral RNA Mini Kit (Qiagen, CA-EUA), followed by a RT-PCR to amplify a small portion of 1639 nucleotides entailing the Envelope (ENV)/NS1 region (1467-3106 according to AF489932 reference sequence) of the viral genome, employing primers pair 3A-4B, described elsewhere [29]. For this purpose we used the QIAGEN OneStep RT-PCR Kit (Qiagen, CA-EUA) according to manufacturer’s instructions, and. Thermocycling conditions consisted of a single step of 50°C for 30 minutes for reverse transcription and 95°C for 15 minutes for reverse transcriptase inactivation, followed by 40 cycles of denaturation at 94°C (30 seconds), annealing at 63°C (60 seconds), extension at 72°C (2 minutes) and a final extension at 72°C (10 minutes). PCR products were confirmed by 1.5% agarose gel electrophoresis stained with SYBR Safe DNA gel stain (Invitrogen) and visualized under blue light. Subsequently, PCR specific bands were sliced from the gel and purified with the Qiagen Gel Extraction Kit (Qiagen, CA-EUA), following manufacturer’s instructions. Quantification of DNA amplicons was carried on with Qubit® fluorometer (Life Technologies) and finally sequenced by Sanger-based technique employing fluorescent dideoxynucleotides at the DNA sequencing Platform at Fiocruz Foundation, Rio de Janeiro.

### Phylogenetic analyses

The obtained sequences (GenBank accession numbers MK972823 and MK972824) were manually edited and aligned with BioEdit v7.2.5.0 software using different DENV-2 reference sequences representative for the different genotypes and available at GenBank (accession numbers can be consulted in the Supplementary Information 1). Alignment of the final 378 sequences is available from the authors upon request. Evolutionary model was obtained with JModeltest v2.1.10 software (GTR+I+G) [30], chosen according the Akaike Information Criterion (AIC) and the Maximum Likelihood value (lnL), and phylogenetic trees were constructed using the Maximum likelihood method with RAxML v7.0 software. The robustness of the phylogenetic grouping was evaluated by bootstrap analysis with 1000 replicates. Phylogenetic tree was finally visualized in Figtree v1.4.3 software (available at http://tree.bio.ed.ac.uk/software/figtree/).

### Single nucleotide polymorphisms assessment

The presence of single nucleotide polymorphisms (SNP) of potential clinical and epidemiological relevance at ENV/NS1 regions was manually determined by comparing the nucleotide and amino acid changes in the generated sequences with the remaining sequences from As/Am genotype, employing as references Brazilian sequences obtained from the states of Rio de Janeiro and São Paulo, during 2007-2011 (GenBank accession numbers: JX286516-JX286521, JX286550/51, and MN589858-MN589884). The translation of the nucleotide sequences to amino acids was performed employing the BioEdit v7.2.5.0 software.

### Spatiotemporal dispersion analyses

Sequences generated within this work plus other 135 DENV-2 sequences obtained from the GenBank (Supplementary Information 1), also employed for phylogenetic analysis, were used, together with their corresponding epidemiological data, to carry on further temporal and geographical estimations of the evolutionary process. First, the presence of sufficient temporal signal in the dataset was evaluated with the TempEst v1.5.1 software [31], to proceed then with phylogenetic molecular clock analysis. The time scale of the evolutionary process was assessed employing the sequences collection dates obtained from the GenBank annotations, and a GTR+I+G4 nucleotide substitution model, a relaxed uncorrelated lognormal molecular clock model, and a Bayesian Skyline coalescent tree prior. Migration events and their most reliable migration pathways were estimated using a non-reversible discrete phylogeography model with the Bayesian stochastic search variable selection approach, and a continuous-time Markov chain (CTMC) migration rate reference prior. The analysis was implemented in BEAST v1.8.1 software package [32]. The Markov Chain Monte Carlo analysis was run for 100 million generations and convergence was assessed in Tracer v1.7 (http://tree.bio.ed.ac.uk/software/tracer): 10% of the sampling trees were discarded as burn-in and accepted effective sample sizes (ESS) values should be higher than 200. The 95% highest posterior density (HPD95%) interval was considered to estimate uncertainty for each estimated parameter. A maximum clade credibility tree (MCCT) was constructed with the TreeAnnotator v1.8.1 software (part of the BEAST package) after discarding 10% of the sampling, and visualized with the FigTree v1.4.3 program (available at http://tree.bio.ed.ac.uk/soft-ware/figtree/).

## Results and discussion

All of the ten analyzed samples tested positive for DENV-2 through the RT-PCR under Lanciotti’s protocol. In general, they all belonged to adults’ patients between 23 and 61 years old, being 6 males and 4 females. None of them presented clinical signs of severity. Their respective epidemiological characteristics can be found as Supplementary information 2. Two out of this ten samples were selected for DENV-2 genetic sequencing based on their epidemiological characteristics: sample number 80, belonging to a 38 years-old female patient from the district of Vassouras who manifested the infection in mid-January, and sample number 230, belonging to a 37 years-old male patient from the district of Volta Redonda manifesting dengue fever in mid-March.

Phylogenetic analysis performed over the two generated sequences showed that DENV-2 strain detected in both patients sera belongs to the same lineage of As/Am genotype that caused the outbreak in 2008. However, it grouped in a totally separated cluster from sequences previously obtained from different locations of Brazil (Figure 1). The current lineage has been previously described by Mir et al. as lineage IV, which dominated the epidemics in South and Central America during the 2000s [33].

**Figure 1.**
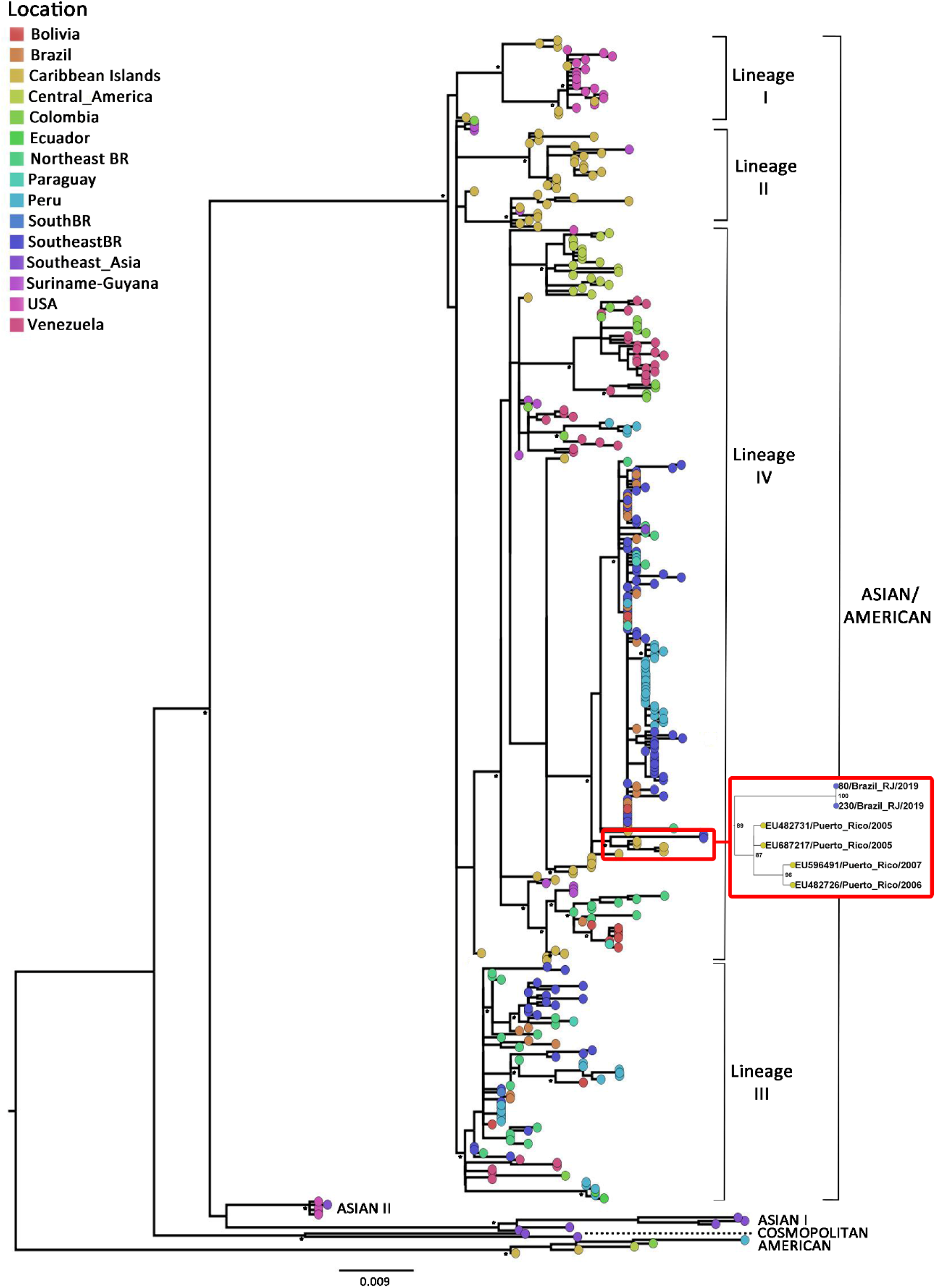
Maximum likelihood phylogenetic tree of DENV-2 partial envelope and NS1 coding sequences. DENV-2 dataset includes partial sequences (positions 1482-3107) principally from the As/Am (n=361), American (n=5, from Colombia, Peru, El Salvador and Caribbean Islands), Asian I (n=5, from Southeast Asia [Thailand and Cambodia]), Asian II (n=5, from the United States of America [USA] and Southeast Asia [Papua New Guinea]) and Cosmopolitan (n=2, from Southeast Asia [India and Singapore]) genotypes. As/Am genotype includes sequences from USA (n=19), Central America [Mexico (n=5), Nicaragua (n=13), Guatemala (n=1), Belize (n=1), El Salvador (n=1)], Guyana (n=1), Suriname (n=11), Ecuador (n=2), Venezuela (n=35), Colombia (n=15), Peru (n=48), Bolivia (n=12), Paraguay (n=5), Brazil [BR] (n=137, which includes 77 from the southeast region and 32 from the northeast) and from the Caribbean Islands which includes Jamaica (n=1), Puerto Rico (n=13), Cuba (n=2), Dominican Republic (n=4) and the Lesser Antilles islands (n=36) conformed by Trinidad and Tobago, the Virgin Islands, Aruba, Dominica, Barbados, Grenada, Martinique, among others. Four main lineages (I to IV) are identified within As/Am genotype. Taxa are represented with circles and colored according to the geographic origin of each sequence as indicated at the legend (up left). The small cluster containing isolates from the state of Rio de Janeiro in 2019 is zoomed-in, and the two sequences generated in this study are indicated with names 80/Brazil_RJ/2019 and 230/Brazil_RJ/2019. The asterisk in the nodes corresponds to bootstrap values higher than 70, obtained with 1000 replicates. The scale bar indicates the genetic distances. The branch lengths are drawn to scale with the bar at the bottom indicating nucleotide substitutions per site.

Surprisingly, this year’s sequences grouped together with sequences from Puerto Rico of 2005-2007, in a highly supported cluster (Figure 1). These findings suggest that a new viral strain may have been introduced into the state of Rio de Janeiro, which in effect, is genetically different from viral strains that kept on circulating basally in other regions of the country, at least in the viral covered region under study.

Based on phylogenetic results, we proceeded to identify the genetic positions on viral sequences involved in genetic distance between the viral isolates under study and the remaining Brazilian ones from lineage IV. Twenty-four different SNPs were detected, being 14 on the partial sequenced ENV region (genetic positions 1482-2421) and 10 on the partial NS1 (genetic positions 2422-3107) (Supplementary information 3). Only five of them were non-synonymous, provoking an amino acid change on the viral proteins: A2023G+G2024C-S643A and C2276T-A727V on ENV; A2434G-I780V and C2878A-L928M on NS1. To notice, simultaneous nucleotide substitutions on positions 2023 and 2024 were detected exclusively in the cluster conformed by sequences generated in this work and from Puerto Rico. Furthermore, substitution C2878A-L928M was encountered only in our sequences and C2276T-A727V only in one sequence within lineage III from As/Am genotype, while A2434G-I780V was widely detected among sequences within lineage IV, and other As/Am lineages and genotypes as well. However, it was absent in the ones representing Brazilian epidemic strain from 2008. No known effect has been found in literature for any of these substitutions. On the other hand, among the 19 synonymous substitutions, three of them were unique for our sequences: G1710T, A1995G and C2307T, located within the ENV. Conversely, from the remaining 16, only half was detected in sequences from lineage IV. However, none of them among the ones from the 2008 strain. The other half was detected in sequences of other As/Am lineages and/or genotypes (Supplementary information 3). In short, all these observations lead us to suspect that the viral strain circulating currently in the state of Rio de Janeiro may have indeed, a different origin than the one that caused the outbreak in 2008 and spread to other regions of the country, mainly through the Southeast.

Another noteworthy fact is that 2019 cases presented different clinical-epidemiological characteristics in contrast with 2008’s. During the first semester of 2008, 209,309 cases were confirmed, including 10947 hospitalizations and 230 deceases, affecting mainly the age group of children under 15 years old [34]. During the same period by 2019, 27,913 probable dengue cases were notified in the state of Rio de Janeiro, of which 59.6% were confirmed clinical and laboratory. Only 267 of them required hospitalization, and no deceases have been yet reported. Most cases occurred in young adults between 20 and 49 years old, with no clear predominance of any gender [35]. Likewise, samples analyzed in this study, corresponded to young adults’ patients, both presenting a classic dengue fever (Clinical end epidemiological characteristics are available as Supplementary Information 2). It is important to mention that also during 2019, a chikungunya outbreak of great magnitude is scourging the state of Rio de Janeiro [35]. It could be likely, that vector competence by both viruses may be leading to a lower circulation of DENV-2, and thus, a reduced number of cases. This scenario may be influencing the marked contrast between the total number of cases.

Considering the phylogenetic results, and to have a more detailed picture of where this new DENV-2 strain comes from, we conducted temporal and geographical analysis with a reduced dataset of 137 sequences conforming lineage IV group of As/Am genotype. Sequences within this cluster are indicated in the Supplementary Information 1. They exhibited a positive correlation between genetic divergence and sampling time (R^2^= 0.6824) and appeared to be suitable for phylogenetic molecular clock analysis, which was after that, implemented in BEAST package [32], as described in the above section. These results suggest that the viral strain detected in our study originated in Puerto Rico (Posterior state probability [PSP]=0.81) in 2002 (95% HPD: 2000–2004) (Figure 2), and was then introduced in our state, where it started to spread in late 2018 (95% HPD: 2018–2019). To notice, no DENV-2 case has been diagnosed during the course of 2019 in the Caribbean islands [36], which clearly suggests that Rio de Janeiro’s 2019 strain could have been circulating elsewhere, up to its arrival into the state, or could have been replicating silently for some time up to rise to a detectable level. It would be likely that actually, the viral strain that entered into the state of Rio de Janeiro might well be introduced first into a different Brazilian state and spread then to Rio de Janeiro, based on the increasing DENV-2 reported cases of other states. As previously mentioned, since 2017, DENV-2 cases have been increasing and spreading through the central-west region, reaching the southeast region in 2018 and early 2019. The latter was responsible for 65.7% of the cases reported until March 2019 and is witnessing local epidemics in several municipalities of three of the four states that integrate it: São Paulo, Minas Gerais, and Espírito Santo; coincidentally, the three states that surround geographically the fourth integrant of the group, the state of Rio de Janeiro [26]. Nevertheless, no molecular epidemiology analyses have been carried on to study genetic viral characteristics on those cases, and to confirm this statement we will need further sequencing yet unavailable. In fact, important caveats should be taken into account for the posed hypothesis: the limited number of available sequences of Latin America covering the region under study, and the lack of Brazilian sequences from this last 5 years period. These two aspects, if different, could be responsible for dissimilar estimations. The lack of DENV-2 Brazilian sequences over recent times goes hand-in-hand with the low circulation of this serotype across the country. Nevertheless, filling this information gap would be determinant to define when this new strain has actually entered into our territory.

**Figure 2.**
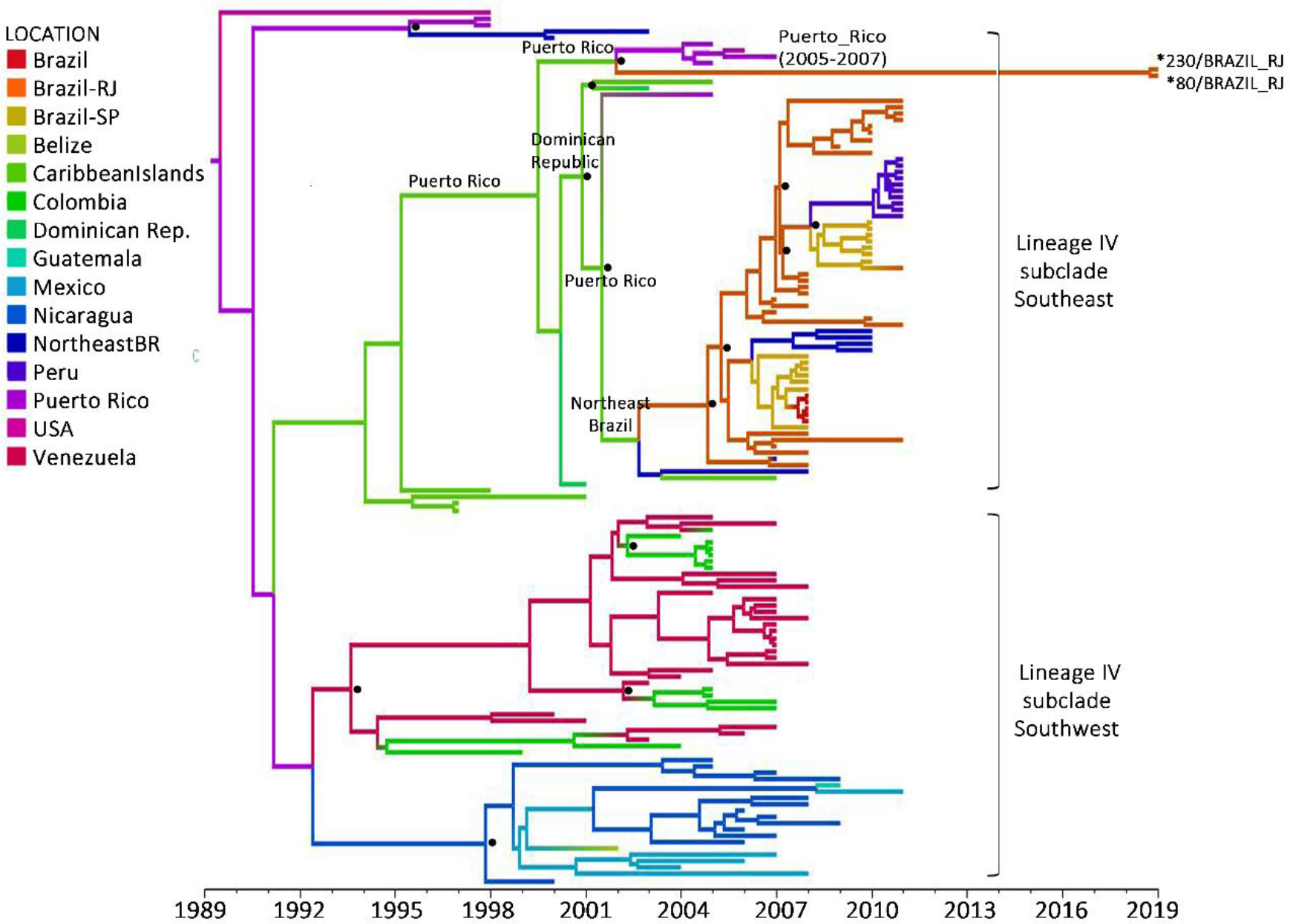
Time-scaled Bayesian Maximum Clade Credibility tree corresponding to the lineage IV of As/Am genotype. Two major highly supported subclades are identified: southeast and southwest. Brazilian sequences are all included in the southeast subclade, and isolates under study are highlighted with an asterisk (*). Branches are colored according to the most probable location (legend shown on the left side) of their parental node inferred by discrete phylogeographical analysis. Location at key nodes involved in the introduction of subclade southeast into Brazil are highlighted. Posterior state probability values higher than 0.7 at key nodes are represented by a dot. All horizontal branch lengths are drawn to a scale of years. Time scale can be observed in the x-axis. The tree is automatically rooted under the assumption of a relaxed molecular clock. In the color-legend, Caribbean Islands includes the countries of Jamaica, Cuba, Virgin Islands, Martinique and Saint Kitts and Nevis.

Even though it is not the main focus of this note, the spatial and temporal origin of lineage IV estimated in our analysis is consistent with results obtained by other groups [15, 33]. According to our calculations, this lineage probably arose in Puerto Rico in middle 1989 (95% HPD=1985–1993) and became the dominant lineage in South and Central America from the early/middle 2000s onwards. The subclade that spread through the southeast pathway, probably arose with the introduction of a virus from Puerto Rico (PSP=0.67) into the northeast region of Brazil around 2003 (95% HPD=2001-2004), moving then to the state of Rio de Janeiro (PSP=0.65) in 2005 (95% HPD=2004-2006), and spreading thereafter through the southeast region. This northeast passage, however, presented a low probability (PSP=0.3). Drumond and collaborators discussed in their work this possible migration pathway through the north as well. Nevertheless, their observation was statistically unsupported [15]. For Mir and collaborators, however, the circulation of this Southeast subclade, named in their work IV-SA4, started after the introduction of the virus from the Great Antilles directly into the southeast Brazilian region in 2004, moving then to the northeastern region, as well as to other countries in South America [33]. The similarities detected between our work and those already mentioned, give extra credibility to the estimates made about this new viral strain detected in the state of Rio de Janeiro.

According to our estimations, the evolutionary rate of lineage IV was 9.5×10^−4^ substitutions/site/year (95% HPD=7.7×10^−4^–1.2×10^−3^ substitutions/site/year), exactly the same reported by Mir et al [33]. To notice, the strain giving rise to the virus isolates detected this year in the state of Rio de Janeiro, presented a nucleotide substitution rate at the lower limit of the 95% HPD interval, which would be consistent with slower evolution dynamics and a delayed detection.

On the other hand, our results regarding DENV-2 population dynamic of lineage IV showed a clear drop in population size between 2005 and 2010, and remained steady until this year (Figure 3). This analysis is representing the lineage IV as a whole, including not only the Brazilian strains but the remaining from Latin America too. The dropping and posterior steady demographic behavior of lineage IV could be consistent with the fact that other DENV serotypes and arboviruses like ZIKV and CHIKV have been responsible for the main epidemics of these last years in the American continent [37–39]. Furthermore, the four DENV serotypes are circulating in the Americas, and have been circulating simultaneously in several countries [40]. Even though it may be difficult to trace the exact history of each DENV serotype in the whole Latin America, it is visible that more than 55% of the total dengue cases notified by the Pan-American Health Organization (PAHO) belonged to Brazil [41]. This means that in a certain way, Latin American DENV history is capturing the Brazilian one. In this regard, and combined with epidemiological data [37, 41–42], it can be suspected that beyond particular places, DENV-2 lineage IV has not been predominant in the Americas since around 2000, which potentially reflects on its demographic reconstruction dynamics.

**Figure 3.**
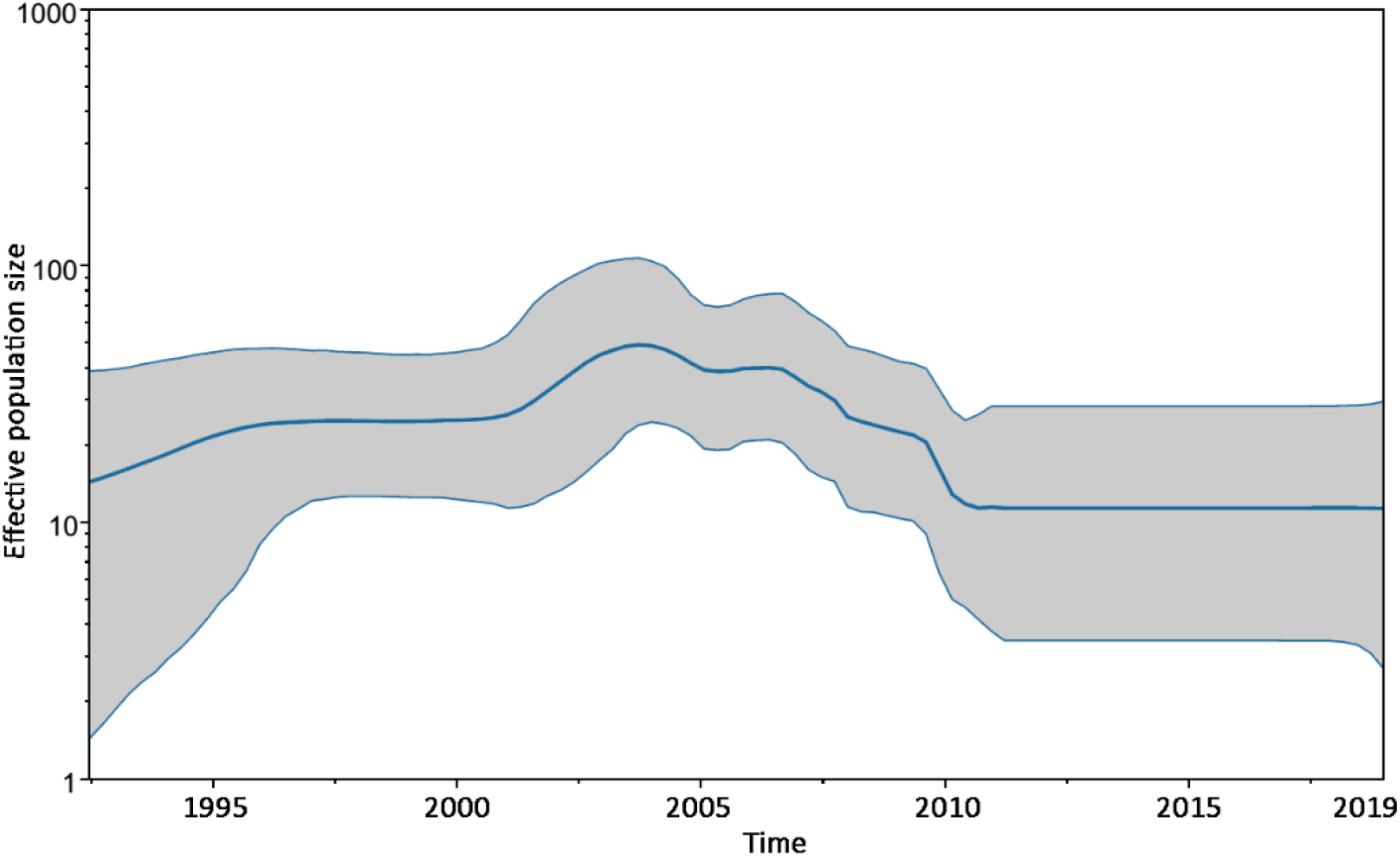
Demographic reconstruction of lineage IV of As/Am genotype. Changes in effective population size since the time of the most recent common ancestor were estimated using the uncorrelated lognormal relaxed molecular clock and the Bayesian Skyline coalescent model. Middle blue line represented the mean value for the effective sample size, while grey areas the 95% confidence interval.

The phenomenon of DENV-2 lineage replacement across successive epidemic outbreaks has been well demonstrated by Mir and collaborators [33]. It should be considered that the circulation of lineage III (that dominated the Caribbean and South America in the 1990s) and IV in the state of Rio de Janeiro, was separated by an 8-year period. This might mean that, aside from the potential viral differences between both lineages itself [16], a generation of children born during this inter-lineage silent period had never be exposed to this serotype, by the time lineage IV entered the state. This fact could be somehow involved in the changes observed in 2007-2008 outbreak regarding the age-group affected.

In effect, a recent PAHO web bulletin is warning Latin American countries about a particular characteristic of the current dengue epidemic, which is threatening not only Brazil but the whole region too: DENV-2 seems to be affecting mostly children and adolescents under the age of 15. In Guatemala, they represent 52% of total cases of severe dengue, while in Honduras, they constitute 66% of all confirmed deaths. Probably, this may be due to their lack of immunity, a consequence of their shorter age and less exposure to the virus in the past [43]. This behavior has not yet been observed in Brazil. However, the current evidence and the learnings from the 2008 outbreak give clear indications that surveillance and control of the current epidemic need to be strengthened.

The facts provided by our work combined with the remaining pieces of evidence, seek to contribute with the health department and medical workers in the outbreak response, emphasizing the necessity of active disease surveillance, including laboratory diagnosis, vector control, and healthcare professionals training for appropriate clinical diagnosis and clinical management of patients with dengue. Working together, we may then be able to ensure a response against the current DENV-2 epidemic scenario, attempting to prevent the outbreak from being of significant magnitude like the one the state of Rio de Janeiro has already experienced a decade ago.

